# Lifestyle and transcriptional signatures associated with ethnicity/race-related variations in the functional connectome

**DOI:** 10.1101/2025.08.27.672637

**Authors:** Ziteng Han, Kexin Wang, Tiantian Liu, Yunxiao Ma, Guoyuan Yang, Tianyi Yan

## Abstract

Understanding the variation of functional architecture across individuals and populations is fundamental to advancing our knowledge of human health and behaviour. Yet, while functional organization differences related to ethnicity/race are consistently reported, their underlying mechanisms remain poorly understood. Here, we apply precision individualized functional mapping to systematically investigate ethnicity/race-related differences in the brain’s intrinsic organization and their associations with lifestyle and transcriptional signatures. We show that variations in network topography and functional connectivity across ethnic/racial groups follow a hierarchical sensorimotor– association axis and are constrained by brain morphology. Importantly, we identify lifestyle factors— particularly education and substance use—that significantly mediate these associations between ethnicity/race and functional connectivity. Leveraging human brain gene expression data, we further demonstrate that cortical transcriptional patterns are spatially aligned with ethnicity/race-related variability in functional connectivity. Gene ontology analyses of associated genes reveal significant enrichment in biological processes such as synaptic signalling and neuronal system development. Together, these findings uncover a multi-layered framework linking ethnicity/race-related differences in brain function to structural constraints, lifestyle influences, and molecular signatures, and advance a more comprehensive and equitable understanding of human brain diversity.

## Introduction

As neuroscience strives for greater generalizability, applicability and equity, increasing attention has turned to ethnicity/race-related differences in brain structure and function[1, 2]. These differences are intertwined with variations in human behaviours and psychiatric outcomes[3, 4]. Neuroimaging evidence has consistently shown that that anatomical properties such as cortical thickness[5], curvature[6] and gray matter volume[7, 8] vary significantly across ethnic/racial populations.

Beyond structural variation, differences have also been observed in intrinsic cortical activity[9] and connectivity profiles[10, 11]. Precision mapping techniques further reveal that ethnicity/race-related variability in functional network topography is heterogeneously distributed across the cortical mantle[11], aligning with fundamental properties of brain[12]. However, given the complex interplay and decoupling of brain structure and function[13, 14], studies directly examining the associations between trans-population differences in brain structure and function are lacking. The extent to which ethnicity/race-related differences in the intrinsic functional organization can be explained by differences in cortical morphometric features remains to be systematically investigated.

Ethnicity/race is a social construct embedded within cultural, environmental, and genetic contexts[15]. Focusing narrowly on superficial group differences may risks oversimplification, potential bias, and inequitable treatment of specific populations. Instead, a deeper understanding requires identifying the root causes of ethnicity/race-related differences[3, 5, 16, 17]. Prior studies have shown that socioeconomic status strongly influences ethnicity/race differences in health outcomes (e.g., mortality) and treatment-seeking behaviours[18–20], and significantly impact the development of brain structures[21] and neurocognitive abilities[22]. Moreover, exposure to social stressors and negative experiences can contribute to ethnicity/race-related variability in neurology and psychopathology (e.g., ageing and threat response)[5, 23, 24]. Recent epidemiological evidence highlights that unhealthy behavioural factors, such as alcohol dependence and lack of physical activity, pose additional risks for both mental and physical health[25, 26], and can explain ethnicity/race-related health inequities[16]. Nevertheless, it remains poorly understood whether and how comprehensive lifestyle mediates brain functional variability across populations, a gap that hinders our understanding of the origins of ethnicity/race-related variations in human brain organization.

Understanding the molecular correlates of brain-wide phenotypes in vivo is important in neuroscience[27, 28]. The Allen Human Brain Atlas (AHBA), which provides a brain-wide map of transcriptional activity across thousands of genes, has been instrumental in linking gene expression to neurodevelopment[29], cortical functional hierarchy[30], psychiatric disorders[31, 32], and inter-individual variability[33]. These findings provide valuable insights into the microscale mechanisms that underlie relevant brain circuits. Importantly, recent transcription studies of postmortem brain tissues from African American (AA) individuals have revealed that genetic ancestry influences gene expression in the brain, with both genetic variations and environments contributing to ancestry-associated expression changes[34]. Ancestry-associated differentially expressed genes (DEGs) are primarily enriched in immune-related pathways[34, 35] and linked to the heritability of neurological disorders[34], echoing the observed prevalence differences across populations[36–38]. Despite these advances, it remains largely unknown how ethnicity/race-related multifaceted factors modulate gene expression and the transcriptomic mechanisms underlying ethnicity/race-related variability in brain functional organization.

To address these gaps, our study investigates the mechanisms underlying ethnicity/race effects on brain functional connectome from the perspectives of anatomical constraints, lifestyle and environments, and transcription. We leveraged multimodal datasets from the Human Connectome Project (HCP), including the HCP-Young Adult (HCP-YA) and HCP-Development (HCP-D) cohorts, which provide diverse ethnic/racial group data from the U.S. population. Specifically, we **first** utilized the advanced multisession hierarchical Bayesian model (MS-HBM) to delineate the individual-specific functional topography and functional connectome architecture. We then used brain-based predictive modelling and linear regression to demonstrate that ethnicity/race is associated with inter-individual variability in resting-state functional connectivity (RSFC) and brain morphological features. **Second**, we revealed shared predictive network features across ethnicity/race and various lifestyle variables. We also demonstrated that the functional connectivity correlates of ethnicity/race are partially mediated by lifestyle factors. **Finally**, by aligning gene expression patterns with functional connectivity features, we revealed transcriptional mechanisms underlying ethnicity/race variations in brain functional organization. Taken together, our work adheres to the latest theoretical framework for neuroimaging research related to ethnicity/race[15] and provides an integrative framework for understanding how structural, lifestyle, and molecular factors converge to shape population-level variability in the brain function.

## Results

We employed two independent large-scale datasets, the HCP-YA and HCP-D datasets, and focused on AA and white American (WA) participants. Quality control process followed a previous study[39]. Afterwards, 101 AA participants and 721 WA participants from the HCP-YA dataset (*N* = 822; age, 22 to 37 years; 435 females) and 68 AA participants and 404 WA participants from the HCP-D dataset (*N* = 472; age, 6 to 21 years; 253 females) were included for subsequent analyses. Ethnic/racial groups were categorized based on self-reported information.

### Ethnicity/race-related variability in brain functional topography and connectome

The MS-HBM, which has been shown to capture the functional organization of individuals precisely, was adopted to generate individual-specific cortical parcellations[40]. In the MS-HBM framework, each cortical region is assumed to exhibit a unique RSFC pattern, and a unified statistical framework is established that accounts for inter-individual RSFC variability, intra-individual RSFC variability and spatial constraints (i.e., encouraging adjacent brain locations with slight changes in RSFC to be assigned the same label) to delineate individualized boundaries under the guidance of group-level parcellation (Schaefer400)[41]. While previous research has characterized individual variability at the level of network topology[42, 43], the MS-HBM offers a deeper level of insight by identifying the fundamental functional units that form these networks within each individual. Resting-state functional magnetic resonance imaging (rs-fMRI) data from all valid sessions of each participant were entered as inputs of MS-HBM.

To investigate group-level differences in functional topography, we randomly selected 40 AA and 40 WA demographically matched individuals. Using their individual-level parcellations derived from the MS-HBM, we obtained group-level AA400 and WA400 parcellation maps via a maximum probability fusion approach[44] (**Supplementary Fig. 1**). Similarity analyses, quantified with the Dice coefficient, revealed that parcellation maps from the same ethnicity/race group across different datasets exhibited the highest similarity (**Supplementary Fig. 1**; e.g., AA400 from the HCP-YA dataset compared with AA400 from the HCP-D dataset: 0.844). By contrast, parcellation maps spanning both datasets and ethnicity/race groups exhibited the greatest spatial distinctions (e.g., AA400 from the HCP-YA dataset compared with WA400 from the HCP-D dataset: 0.784). We then assessed the influence of ethnicity/race on individual-level functional topography. Specifically, the similarity of individual-specific parcellation maps between all AA–WA pairs (101×721 in the HCP-YA dataset and 68×404 in the HCP-D dataset) was calculated and averaged. We observed a non-uniform distribution of variability along the cortex, with the greatest variability in heteromodal association cortices (**Fig. 1a and Supplementary Fig. 2**; e.g., control and dorsal attention networks), and the highest similarity in unimodal sensorimotor cortices (especially in the somatomotor and auditory networks).

**Fig. 1.**
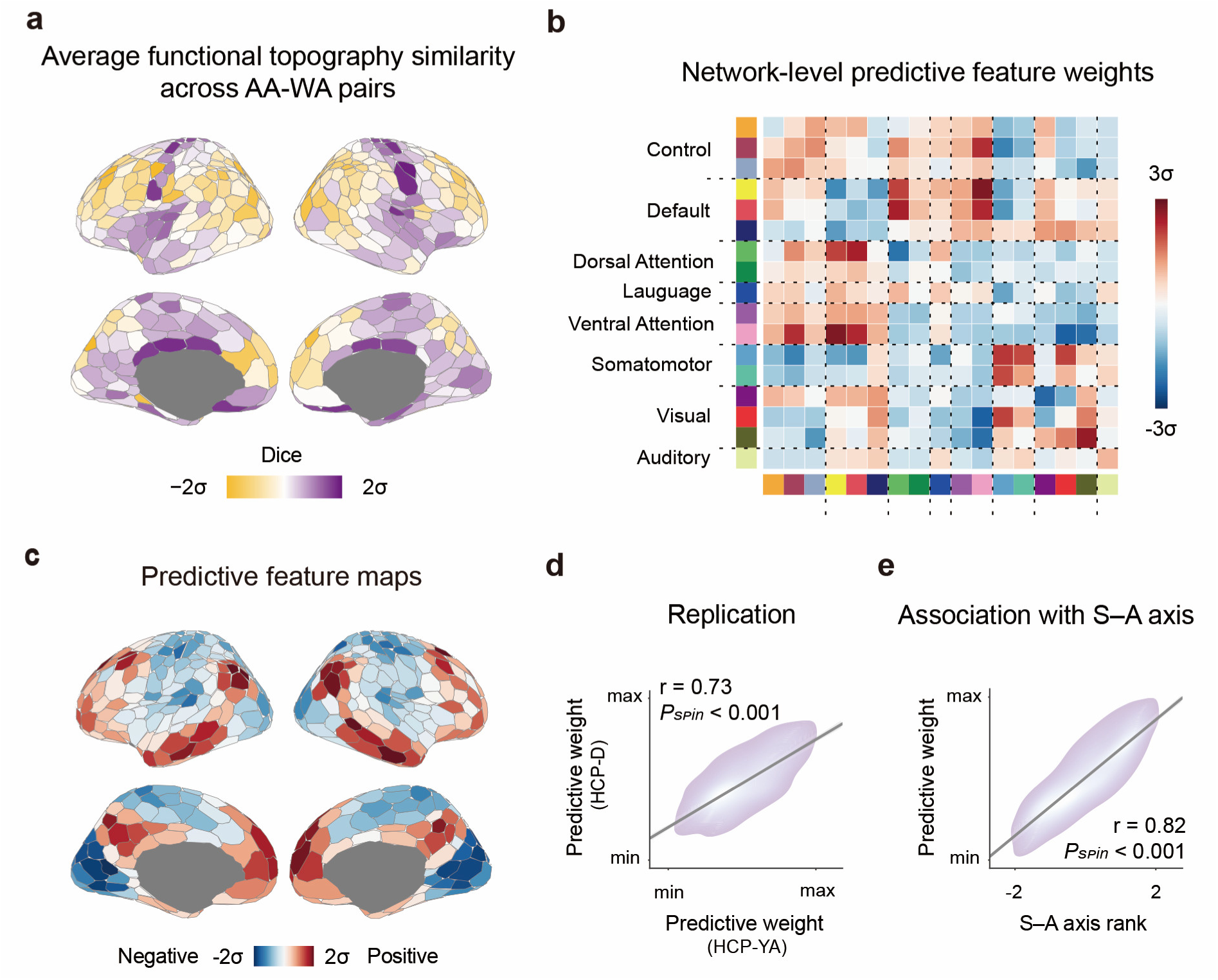
Functional topography and functional connectivity are associated with ethnicity/race. **a**, The average similarity of individual-specific parcellations across all the AA–WA pairs revealed that ethnicity/race-related topographic variability was greatest in heteromodal association cortices and lowest in the unimodal sensorimotor cortices. Brain maps were generated using the Schaefer400 parcellations[41] and visualized with the R package ggseg[48]. **b**, Kernel ridge regression (KRR) models were trained using resting-state functional connectivity (RSFC), while controlling for age, sex, root-mean-square framewise displacement (RMS), intracranial volume (ICV), education and income to predict individuals’ self-reported ethnicity/race. Haufe-transformed regional pairwise feature weights were averaged to the network level based on the Yeo networks. Connections between default and attention networks, as well as within default networks and sensorimotor networks, contributed most to the prediction in both datasets. For visualization, actual values were subjected to *Z* score standardization. **c,** Haufe-transformed weights of each brain region were averaged to generate ethnicity/race-predictive weight maps. **d,** RSFC-based ethnicity/race prediction was performed in both the discovery HCP-YA dataset and the replication HCP-D dataset. Predictive weights were highly reproducible across the two independent datasets (*r* = 0.73, *Pspin* < 0.001). **e,** Ethnicity/race-predictive weight distributions were significantly correlated with the S–A axis ranks (*r* = 0.82, *Pspin* < 0.001).

The individualized parcellations and group-level Schaefer400 parcellations were then applied to rs-fMRI data to generate RSFC matrices (400 × 400) for each participant separately. For each dataset, AA participants were randomly split into 5 folds, with family structure accounted for, and Hungarian matching was employed to select AA–WA pairs matched on age, sex and root-mean-square framewise displacement (RMS)[39]. To minimize sensitivity to particular data split, the above steps were repeated multiple times until 40 distinct AA splits with matched WA splits emerged.

Multivariate kernel ridge regression (KRR) was employed to distinguish between AA and WA individuals, with RSFC matrices as predictors and age, sex, RMS, intracranial volume (ICV), education and income as covariates[39, 45]. Specifically, nested 5-fold cross validation was performed, with all the training folds used to optimize the regularization parameter and the remaining test fold to evaluate model generalizability on unseen data. The above 5-fold cross- validation step was repeated 40 times, and classification accuracy was averaged across folds for each split. In the HCP-YA dataset, our multivariate approach achieved high classification performance, with an accuracy of 85.3% ± 2.3% (mean ± standard deviation) using individual-specific parcellations and 77.9% ± 2.6% using Schaefer400 parcellations, both significantly exceeding chance levels (both *P* < 0.001). In the HCP-D dataset, model performance decreased slightly but remained significantly above chance (both *P* < 0.001), with accuracies of 81.7% ± 5.0% for individual-specific parcellations and 62.5% ± 5.1% for Schaefer400 parcellations. These results emphasize the importance of treating individual-specific topographic features as phenotypes and highlight the robust ethnicity/race-related variability in functional topography and functional connectivity patterns.

To interpret the network features learned by the model, the regression models were inverted using Haufe’s transform, yielding prediction feature matrices[46]. **Figs. 1b and Supplementary Fig. 3** show the averaged pairwise region-level feature weights at the network level, indicating that ethnicity/race-associated functional connections constitute a distributed set of large-scale network circuits. The most prominent links were observed between the default and attentional networks, within the default networks, and within the sensorimotor networks for both datasets. Moreover, to quantify regional contributions, average feature weights of cortical regions were calculated (**Fig. 1c and Supplementary Fig. 3**). We found that discriminating features derived from the HCP-YA and HCP-D datasets exhibited high generalizability and significant spatial correlations (**Fig. 1d**; *r* = 0.73, *P_spin_* < 0.001). Across both datasets, contribution weights followed a hierarchical distribution across the cortex, with opposite contributions between heteromodal cortices and unimodal cortices. This hierarchical layout, reflecting diverse neurobiological properties, corresponds to the well-established cortical sensorimotor–association (S–A) axis[47]. Using spatial permutation testing, we confirmed that the contribution of cortical regions to ethnicity/race prediction were aligned with the S–A axis (**Fig. 1e**; *r* = 0.82, *P_spin_* < 0.001).

### Ethnicity/race-related variability in brain functional connectome is constrained by the fundamental properties of brain organization

The spatial and physical patterns of brain anatomy are known to constrain neural dynamics[13]. Thus, we next examined whether ethnicity/race-related variability in functional connectivity could be explained by the corresponding variability in cortical morphology. To this end, we constructed a morphometric similarity network (MSN) for each individual, which integrates multiple structural MRI features (*Z* scores) into a biologically meaningful and behaviourally relevant connectome (400×400)[49]. Correlation weights were averaged across rows and within groups to generate group-level regional MSN values (**Fig. 2a**). To assess ethnicity/race-related variability in brain morphology, linear regression models (LRMs) were applied while controlling for age, sex, RMS, ICV, education level and income[39]. We observed significant ethnicity/race-related variations in morphometric similarity patterns across the cortex (**Fig. 2b and Supplementary Fig. 4**), especially in the insula, prefrontal cortex, cuneus and lateral occipital cortex. Importantly, this variability was highly reproducible across datasets (**Fig. 2c**; *r* = 0.59, *P_spin_* < 0.001). Moreover, the MSN *t*-map was positively associated with the RSFC-based ethnicity/race predictive weights (**Figs. 2d**; *r* = 0.41, *P_spin_* = 0.012), indicating that brain regions contributing more to RSFC-based predictions also exhibited greater ethnicity/race-related variability in cortical morphology.

**Fig. 2.**
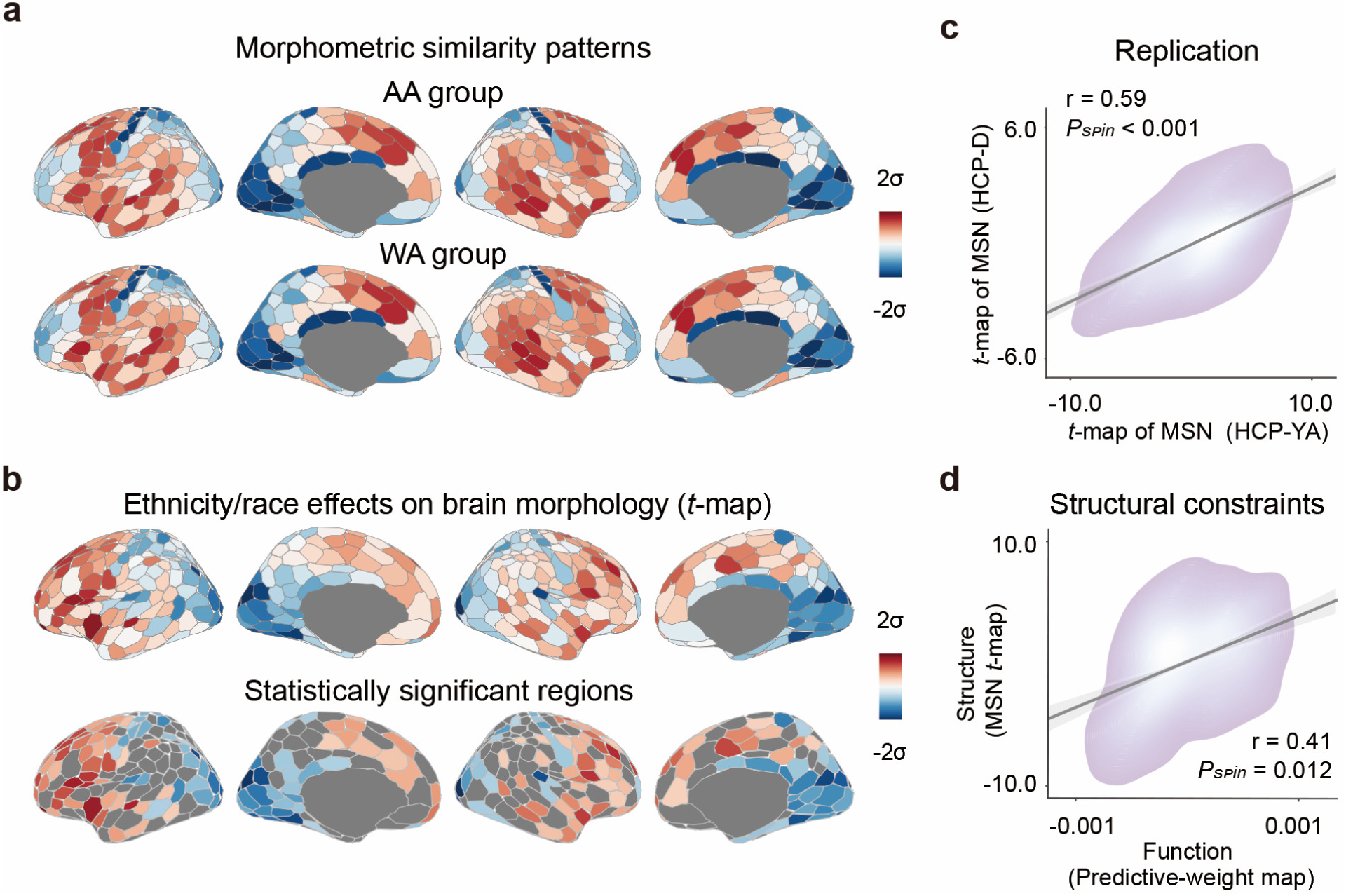
Ethnicity/race-related variability in functional connectivity profiles aligns with ethnicity/race-related variations in cortical morphometry. a, The cortical distribution of mean regional morphometric similarity network (MSN) values was examined, reflecting inter-regional similarity across multiple MRI-derived morphometric features (i.e., cortical thickness, gray matter volume, surface area, Gaussian curvature and mean curvature). Brain maps were generated using the Schaefer400 parcellations[41] and visualized with the R package ggseg[48]. b, Comparison (*t*-map) of regional MSNs across ethnic/racial groups was performed. Linear regression models (LRMs) were fitted to estimate the ethnicity/race-related variations in the MSN, controlling for age, sex, RMS, ICV, education and income. False discovery rate (FDR) correction with *P* < 0.05 for multiple comparisons across 400 regions identified significant ethnicity/race-related effects on cortical morphometry. c, The *t*-map for regional MSN differences was highly reproducible across the two datasets (*r* = 0.59, *Pspin* < 0.001). d, Ethnicity/race-related variability in cortical morphometric similarity accounted for parts of the variance in ethnicity/race-relevant variability in functional connectivity profiles (*r* = 0.41, *Pspin* = 0.012).

Beyond the MSN, we considered other fundamental properties of brain organization. Intracortical myelination, a microstructural marker of neuronal fibres that influences processing efficiency in the nervous system[50, 51], was estimated from the ratio of T1-weighted to T2-weighted images, where higher values reflect greater myelination. Across the cortex, predictive weight maps were inversely related to the cortical myelin content (**Supplementary Fig. 5**; *r* = −0.57, *P_spin_* < 0.001). Together, these results demonstrate that ethnicity/race-related variability in brain functional connectivity is tightly linked to fundamental structural properties of brain organization.

### Lifestyle influences ethnicity/race-related variability in brain functional connectome

Lifestyle factors, including education, income, alcohol consumption, smoking, sleep, physical activity, and social connection, have been shown to influence brain structure and function[52–55]. Having revealed the ethnicity/race-related variability in RSFC architecture, we next sought to investigate the role of these lifestyle factors in explaining this variability. Specifically, our goal was to identify the shared brain-network features that are associated with both ethnicity/race and various lifestyle factors. We selected 6 lifestyle factors (i.e., education, income, substance use, sleep, physical activity, and social relationships) from the HCP-YA dataset and we did not conduct the subsequent analysis for the HCP-D dataset as the majority of lifestyle data were only available for a subset of participants (e.g., those under age 17), which would substantially reduce the AA sample size. We found that several lifestyle factors were significantly associated with ethnicity/race categories in the HCP-YA dataset: education (*r* = −0.15, *P* < 0.001), income (*r* = −0.23, *P* < 0.001), substance use (*r* = −0.08, *P* < 0.05), sleep (*r* = 0.11, *P* < 0.01), physical activity (*r* = −0.18, *P* < 0.001) and social relationships (*r* = −0.08, *P* < 0.05).

Furthermore, all the HCP-YA participants were randomly split into 15 folds, with the constraint that participants from the same family were kept together in the same fold. To minimize sensitivity to any particular data split, this procedure was repeated 40 times. Using KRR models, we showed that all these factors could be predicted significantly above chance from RSFC matrices, with prediction accuracies of 0.19 ± 0.01 for education, 0.11 ± 0.01 for income, 0.19 ± 0.01 for substance use, 0.09 ± 0.01 for sleep, 0.08 ± 0.01 for physical activity and 0.07 ± 0.01 for social relationships (**Fig. 3a ;** Pearson’s coefficient was employed as a prediction performance metric). Moreover, we examined the similarity of Haufe-transformed predictive-network features underlying ethnicity/race and each lifestyle factor. Strong correlations were observed for education (*r* = −0.55), income (*r* = −0.57), substance use (*r* = −0.34), sleep (*r* = 0.36) and physical activity (*r* = −0.38), but not for social relationships (*r* = 0.05; **Fig. 3b**). Predictive-network feature matrices for all the lifestyle factors are shown in **Supplementary Fig. 6.** Finally, by leveraging data capturing the impacts of neighbourhood-level socioeconomic environments on intrinsic brain activity reported by Sydnor et al.[47], we demonstrated that the observed ethnicity/race-related variability aligns with brain– environment associations across the cortical mantle (**Fig. 3c**; *r* = 0.57, *P_spin_* < 0.001 in the HCP-YA dataset; *r* = 0.41, *P_spin_* = 0.002 in the HCP-D dataset), suggesting ethnicity/race-by-environment interaction effects that shape brain functional organization.

**Fig. 3.**
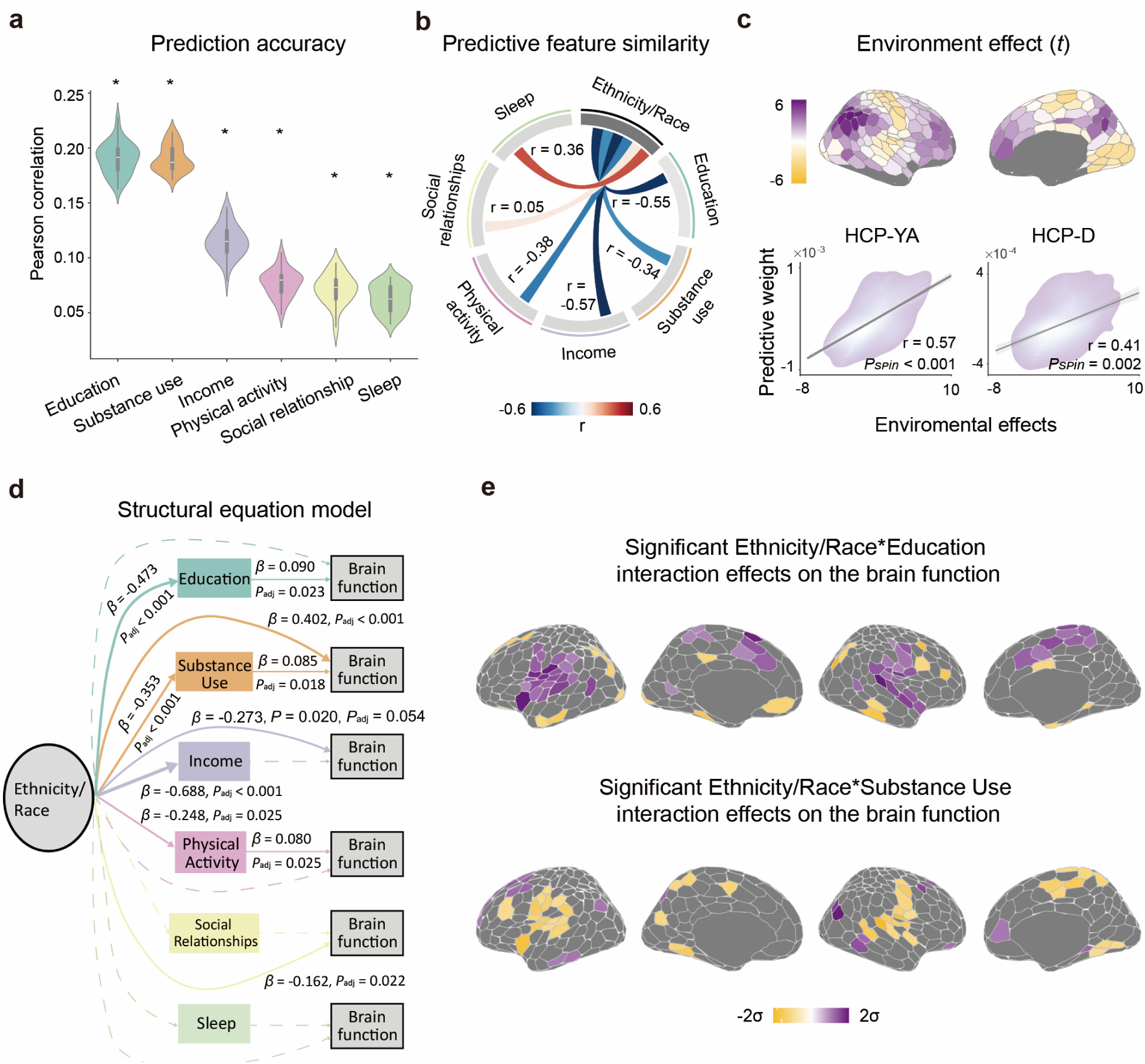
Lifestyle factors mediate the associations between ethnicity/race and brain functional connectivity. **a**, Mean prediction accuracies for lifestyle factors. Asterisks indicate above-chance predictions (*P* < 0.05), determined by comparing true accuracy with 1,000 permuted null models. Violin plots display accuracy distributions across 40 replications. For each violin plot, width represents the density of the accuracy values, and the central box shows the interquartile range and median. **b,** Correlations of predictive feature matrices between ethnicity/race and lifestyle factors, indicating shared brain-network features underlying both ethnicity/race and lifestyle predictions. **c,** Functional connectivity-based ethnicity/race predictive features were significantly associated with effects of neighbourhood socioeconomic conditions on the intrinsic brain activity (HCP-YA dataset: *r* = 0.57, *Pspin* < 0.001; HCP-D dataset: *r* = 0.41, *Pspin* = 0.002). **d,** A structural equation model was fitted for each lifestyle factor, thereby revealing the relationships between ethnicity/race, lifestyle and brain functional organization, while controlling for age, sex, RMS, ICV, education and income. Edges represent regression coefficients (*β* values), with thickness proportional to their absolute magnitude. Significant links (FDR-corrected *P* < 0.05, two-sided) are shown as solid lines; non-significant links as dashed lines. Mediation analysis revealed that education (absolute mediation proportion = 14.57%) and substance use (absolute mediation proportion = 8.07%) significantly mediated the relationship between ethnicity/race and brain functional organization (FDR-corrected *P* < 0.05). **e,** Threshold *β*-maps from linear regression models (LRMs), showing statistically significant (*P*FDR < 0.05) ethnicity/race–education and ethnicity/race–substance use interaction effects on the brain functional organization.

Next, we fitted a structural equation model (SEM) for each lifestyle factor to analyse the relationships among ethnicity/race group, lifestyle and brain functional organization (represented by the principal functional gradient), while controlling for age, sex, RMS, ICV, education and income (**Fig. 3d**). The principal functional gradient captures the dominant patterns of cortical variability in functional connectivity, offering a simplified, low-dimensional representation of brain functional organization in a hierarchical manner[56]. The regression coefficients in the converged SEM suggested that ethnicity/race category was a significant predictor for education (*β* = −0.473, *P*_FDR_ < 0.001), income (*β* = −0.688, *P*_FDR_ < 0.001), substance use (*β* = −0.353, *P*_FDR_ < 0.001) and physical activity (*β* = −0.248, *P*_FDR_ = 0.025). In turn, education (*β* = 0.090, *P*_FDR_ = 0.023), substance use (*β* = 0.085, *P*_FDR_ = 0.018) and physical activity (*β* = 0.080, *P*_FDR_ = 0.025) were significant predictors for brain function, indexed by the first principal gradient of the RSFC profiles. Mediation analyses further revealed that education (mediation proportion = 14.57%, *P*_FDR_ < 0.05) and substance use (mediation proportion = −8.07%, *P*_FDR_ < 0.05) significantly mediated the associations between ethnicity/race category and brain function. Other mediation effects, including income (*P*_FDR_ = 0.784), physical activity (*P*_FDR_ = 0.115), sleep (*P*_FDR_ = 0.140) and social relationships (*P*_FDR_ = 0.673), were not statistically significant after false discovery rate (FDR) correction.

Given the mediating roles of education and substance use, we further investigated how their interactions with ethnicity/race influence brain functional organization across the cortical mantle using LRMs (**Fig. 3e**). We identified 91 cortical regions significantly modulated by ethnicity/race– education interactions, primarily located in the insula, superior temporal, inferior parietal and superior frontal regions. Additionally, 78 cortical regions were significantly influenced by ethnicity/race–substance use interactions, including regions in the insula, superior temporal, rostral middle frontal, and superior frontal cortices.

### Cortical gene expression related to ethnicity/race-induced variability in brain functional connectome

After demonstrating the influence of lifestyle factors on ethnicity/race-related variability in RSFC profiles, we next aimed to identify the transcriptional signatures associated with this variability.

Brain-wide gene expression data were obtained from the AHBA (http://human.brain-map.org)[57, 58] and mapped into 400 region-level parcellations (400 regions × 15,632 genes) derived from the original 3,702 distinct microarray samples. Using partial least squares (PLS) regression[59], an advanced approach for uncovering fundamental relationships between two high-dimensional matrices, we identified weighted cortical gene expression patterns that aligned with the spatial distribution of ethnicity/race predictive features. The first component (PLS1), which explained the largest proportion of variance, accounted for 40.3% (*P_perm_* < 0.001) of the response variance in the HCP-YA dataset and 22.9% (*P_perm_* < 0.001) in the HCP-D dataset.

We found that the PLS1-weighted maps revealed an anterior-to-posterior hierarchy pattern along the cortex, which was remarkably consistent across the two datasets (*r* = 0.99, *P_spin_* < 0.001). Moreover, PLS1 gene expression weights were strongly correlated with the ethnicity/race predictive weights (**Fig. 4a**; *r* = 0.65, *P_spin_* < 0.001), suggesting that genes with higher positive (or negative) weights on PLS1 tend to be overexpressed in regions showing corresponding positive (or negative) ethnicity/race predictive weights. We then ranked the genes based on normalized PLS1 weights and selected those with the top 2,000 positive weights (*Z* > 6.7 in the HCP-YA dataset and *Z* > 4.5 in the HCP-D dataset), denoted as the PLS+ gene sets, as well as the top 2,000 negative weights (*Z* < −6.5 in the HCP-YA dataset and *Z* < −4.6 in the HCP-D dataset), denoted as the PLS-gene sets (**Fig. 4b and Supplementary Fig. 7**).

**Fig. 4.**
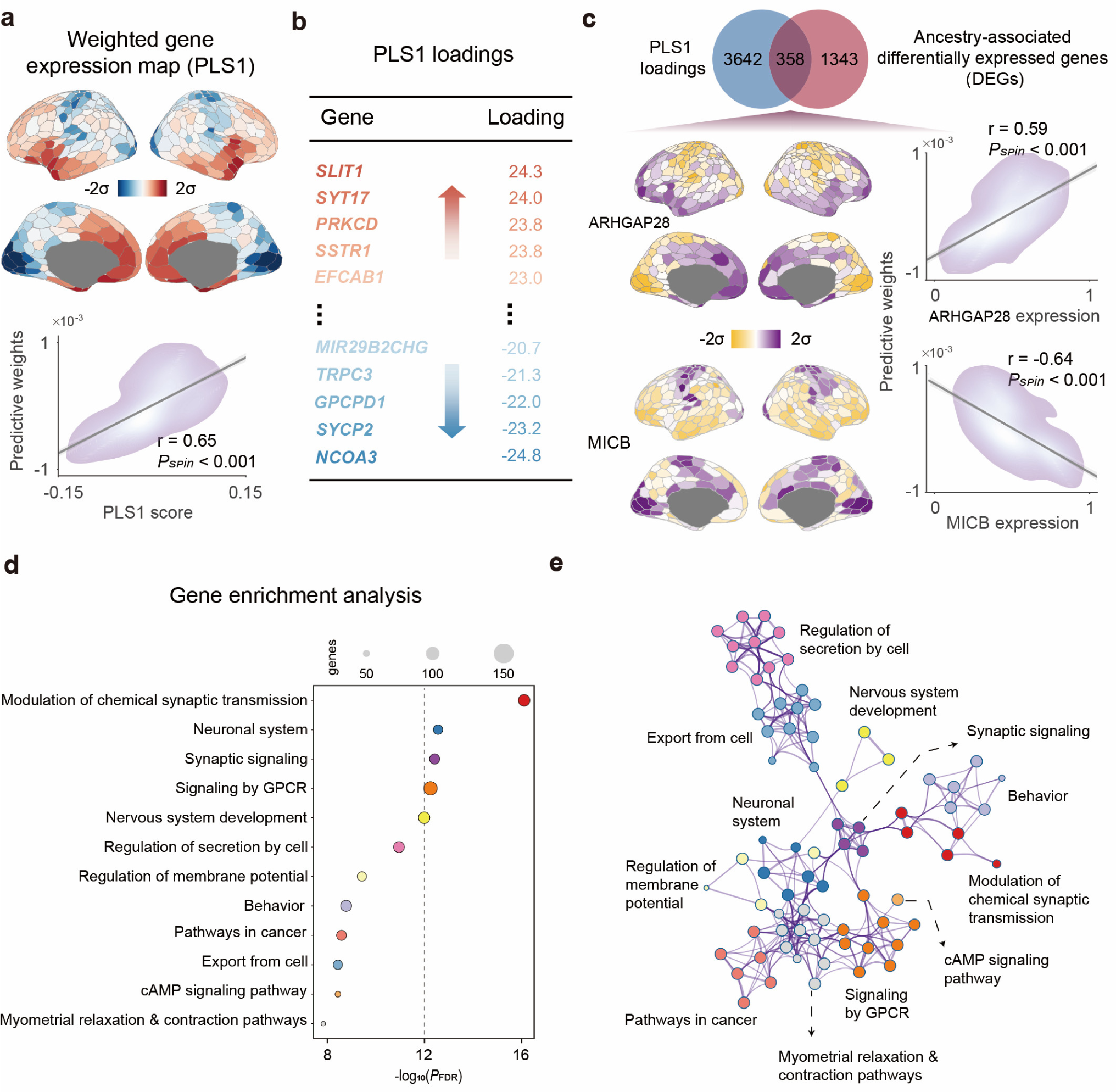
Gene expression profiles associated with ethnicity/race-related variability in functional connectivity. **a**, Weighted cortical gene expression map of the regional PLS1 scores (top panel). Scatterplot showing the relationship between the PLS1 map and the Haufe-transformed predictive weight map (bottom panel, *r* = 0.65, *Pspin* < 0.001). Brain maps were generated using the Schaefer400 parcellations[41] and visualized with the R package ggseg[48]. **b,** Genes were ranked on the basis of their loadings on PLS1. **c,** The PLS+ genes overlapped with ancestry-associated differentially expressed genes (DEGs) reported by Benjamin et al[34]. **d,** Representative enriched terms of the PLS1+ genes, highlighting GO biological processes related to synaptic signalling, as well as Reactome gene sets associated with the nervous system. Circle size denotes the number of genes within a given ontology term, and multi-test FDR-adjusted *P* values are plotted as log10-transformed values. **e,** Metascape network plot of enriched terms, capturing intra-cluster and inter-cluster similarity relationships. Each term is represented by a circle, coloured by cluster identity and scaled proportional to the number of genes involved.

We next conducted an external validation by comparing our PLS1 gene set with the genetic ancestry-associated DEGs identified from RNA sequencing data of postmortem brain tissue[34]. Among the 1,701 DEGs, only 358 (∼21.0%) overlapped with the PLS1 genes in the HCP-YA dataset, 358 (∼21.0%) overlapped in the HCP-D dataset, and 309 (∼18.2%) overlapped in both datasets (**Fig. 4c** <u>and</u> **Supplementary Fig. 7**). Given that ancestry-related gene expression is primarily driven by genetic variations (explaining ∼60% of the changes) rather than the environments (∼15%)[34], this limited overlap between the MRI-derived PLS1 genes and ancestry DEGs suggests that the transcriptional changes associated with self-reported ethnicity/race are largely distinct from those driven by genetic ancestry. One plausible explanation for this distinction is that the MRI-derived PLS1 gene signature is more strongly associated with environmental and lifestyle factors. Delving into the overlapping genes, ARHGAP28, which encodes the Rho GTPase-activating protein[60, 61], exhibited the strongest positive correlation with ethnicity/race predictive weights (**Fig. 4c**; *r* = 0.59, *P_spin_* < 0.001). In contrast, MICB, which encodes ligands for the natural killer cell-activating receptor NKG2D[62], exhibited the strongest negative correlation (**Fig. 4c**; *r* = −0.64, *P_spin_* < 0.001).

To further interpret the PLS- and PLS+ gene sets, we conducted gene enrichment analysis to characterize the enriched biological functions and pathways using Metascape[63]. After employing Benjamini‒Hochberg correction for multiple hypothesis testing and discarding discrete enriched clusters, the remaining significantly enriched terms of the PLS+ genes were rendered into a network (**Figs. 4d–e** <u>and</u> **Supplementary Figs. 7**), including gene ontology (GO) biological processes, such as “modulation of chemical synaptic transmission” and “synaptic signalling”, as well as Reactome gene sets, such as “neuronal system”, “signalling by GPCR” and “nervous system development”. The PLS1-genes were enriched for GO biological processes, such as “metal ion transport” and “potassium ion transmembrane transport”, as well as Reactome pathways, including “neuronal system” and “transport of small molecules” (**Supplementary Fig. 8**). Collectively, these results highlight the distinct functional roles of genes with positive versus negative weights on PLS1.

We further assessed the reproducibility of transcriptional enrichment associated with ethnicity/race-related functional connectivity features. Specifically, we conducted a multi-gene list meta-analysis of PLS1 genes (combining the PLS+ and PLS-genes) across the HCP-YA and HCP-D datasets. The gene lists in these two independent datasets showed substantial overlap, with an odds ratio of 271.88 (**Supplementary Fig. 9,** *P* < 0.001). After statistical correction, the surviving ontological terms were highly shared across both gene lists, and the resulting network demonstrated significant concordance with the discovery enrichment results (**Supplementary Figs. 9**).

## Discussion

In this study, we presented a multi-layered framework for understanding ethnicity/race-related variability in brain functional organization. Using multimodal HCP datasets and precision individualized parcellation, we showed that hierarchical connectivity patterns from sensorimotor to association cortices predict ethnicity/race and align with structural variations across populations.

Furthermore, we observed that the shared brain-network features served as predictors for both ethnicity/race and various lifestyle factors, with education and substance use substantially mediating the associations between ethnicity/race and brain connectivity. We also characterized the transcriptomic architectures underlying the observed ethnicity/race-related variability, thereby providing biological microscale insights into the macroscale properties. Notably, the reproducibility across two large cohorts underscores the robustness of our findings and echoes recent calls for reproducible neuroscience research. Overall, our study integrates structural, behavioural, and molecular dimensions to understand the human brain diversity and contributes to a more generalizable and equitable neuroscience.

Both the spatial configuration of functional regions and the coupling strength between regions are important drivers of variability in functional organization, which is closely linked to non-imaging measures of behavioural and lifestyle measures[64]. Here, we observed that both functional topography and connectivity are associated with individuals’ ethnicity/race. First, spatial variation in functional topography across ethnicity/race groups was greater in heteromodal association cortices than in unimodal sensorimotor cortices. This finding extends previous findings showing that association networks exhibit the greatest variations in functional topography across age windows[65], sex[66], and cultural populations[12]. Second, multivariate patterns of brain-ethnicity/race associations were inversely driven by the connectivity profiles of higher-order association regions versus modality-specific regions, indicating that ethnicity/race-related sociodemographic factors exert widespread and opposite effects on the two poles of the S–A axis[47]. Additionally, ethnicity/race-related differences in functional connectivity profiles were closely aligned with variations in morphometric similarity across groups. This suggests that functional connectivity differences associated with ethnicity/race are constrained by brain morphology, conforming with the well-established concept of cortical structural constraints on intrinsic neuronal activity[13, 67]. These findings further support the homophilic principle of brain organization, which posits that brain regions with similar functional characteristics are more likely to share similarities across multiple biological domains and to be connected via white matter tracts[68].

Given the multidimensional properties of ethnicity/race, a comprehensive perspective is needed to link ethnic/racial diversity with the hierarchical functional organization of the brain[39]. Recent compelling evidence demonstrates that children’s neighbourhood environments exert opposing effects on the fluctuation amplitude of intrinsic activity within the sensorimotor and association cortices[47]. Socioeconomically disadvantageous environments diminish the differentiation of the S– A hierarchy[47]. In this study, we demonstrated that these environmental effects are spatially aligned with ethnicity/race-related variability, suggesting an overlap between ethnicity/race and various socioenvironmental factors. Moreover, it is well-established that environmental and lifestyle factors can shape both brain structure and function[54, 55, 69]. Building on this, our findings reveal shared functional network features underlie both lifestyle factors and ethnicity/race. Specifically, connections between the control and somatomotor networks, between the default and ventral attention networks, and within the somatomotor network were strongly associated with both education level and ethnicity/race. In addition, connections within the default networks, between the default and dorsal attention networks, and between the default and ventral attention networks were robustly predictive for both the substance use experience and ethnicity/race. Indeed, behaviour prediction models may capture broad behaviour dimensions within a given behavioural domain rather than the specific traits they are trained on, such as the *g* factor in the case of cognition and the *p* factor in the case of mental health[45]. Thus, the observed similarity of predictive network features suggests that the ethnicity/race variable and lifestyle factors may be supported by shared underlying meta-factors[70].

Moreover, we found that lifestyle factors—particularly education and substance use—significantly mediated the associations between functional connectivity profiles and ethnicity/race. This is particularly relevant as prior studies have shown that educational attainment and rates of substance use (especially marijuana and alcohol) differ across ethnic/racial populations[16, 71–73]. The principal functional gradient of regions located in the insula, ventral sensory/motor cortex and prefrontal cortex was significantly influenced by the interactions between ethnicity/race and education, as well as by the interactions between ethnicity/race and substance use. These regions serve as crucial control hubs of top-down information processing[74] and overlap with circuits implicated in emotional regulation and threat responses[75, 76]. Moreover, both education and substance use are important components in pathways linking the environment and health status[55]. Educational background is often linked to cognitive reserves[77]. Higher levels of education may enhance brain’s flexibility and adaptability during task processing and decision-making[78, 79], and protect neurogenerative risks[80, 81] in later life. Excessive substance use alters the neurobiology of reward processing, control dynamics and emotion regulation[82–84] and is associated with psychiatric conditions, particularly depression symptoms, and reduced quality of life[85–88].

Together, fully accounting for socioenvironmental disadvantages of specific populations is essential for interpreting neuroimaging-derived differences across populations and for developing more generalizable interventions. Finally, although the lifestyle factors analysed here were relatively coarse, they explained nearly one-quarter of the observed ethnicity/race-related variability, underscoring the need for future work to incorporate more comprehensive measures of lifestyle (e.g., access to healthcare) and neighbourhood-level environment (e.g., population density or percentage of residents living in poverty) to more fully elucidate ethnicity/race-related variability in brain organization and brain–behaviour relationships and to promote equitable translation of neuroscience findings.

After linking functional network properties to transcriptomic data, we identified a set of genes (PLS1) whose weighted cortical expression patterns were significantly correlated with ethnicity/race-related variability in functional connectivity. The PLS1 gene expression map revealed high expression in the primary visual, anterior cingulate, and insular cortex, following the global cortical hierarchy of the brain. Gene-to-function annotation analysis revealed that these genes formed a topologically clustered network enriched for several GO biological processes and Reactome pathways in both the HCP-YA and HCP-D cohorts. The identified GO terms were primarily related to synaptic signalling and the modulation of chemical synaptic transmission, which underpin inter-neuronal communication in the nervous system[89]. This enrichment is consistent with prior reports of population differences in gene expression associated with axon guidance and synaptic signalling pathways[35]. The identified Reactome pathways included the neuronal system and its development, which encompass neuronal progenitor cell proliferation, migration, differentiation, synaptogenesis, and synaptic pruning—all fundamental for establishing a mature and integrated neural system[90, 91]. Importantly, many of these neurodevelopment-related pathways are influenced by environmental and lifestyle factors, such as socioeconomic status[92], alcohol consumption[93], early-life stress[94] and physical activity[95, 96]. Another key Reactome pathway involved G protein-coupled receptor signalling, which regulates cellular responses to hormones, neurotransmitters, and external environmental stimulants[97]. Overall, our findings bridge the gap between ethnicity/race-related variability in brain functional organization and the underlying transcriptional programs, highlighting synaptic and neurodevelopmental pathways as central mechanisms.

Several limitations of the current study should be considered when interpreting our findings. First, we relied on correlation analyses to describe relationships between functional and structural differences across ethnicity/race groups. Future research employing causal modelling techniques is necessary to elucidate the underlying causal architecture of these associations. Second, the generalizability of our findings is constrained by the nature of the datasets used. Our population sample was limited to self-identified WA and AA participants, without accounting for the cultural and geographical factors that may influence the brain. This makes it challenging to generalize our conclusions across the full spectrum of ethnicity/race populations. Third, by using two independent cross-sectional cohorts (HCP-YA and HCP-D) that differ in scanner protocols, we were unable to explicitly model developmental trajectories of ethnicity/race-related differences. Longitudinal datasets, such as the Adolescent Brain Cognitive Development (ABCD) study, are needed to investigate how these brain-wide associations evolve across developmental stages. Fourth, the transcriptome-neuroimaging associations are limited by the small, demographically skewed sample (a sample of six donors—one Hispanic, two AA, three Caucasian—only two of whom provided data for the right hemisphere) of the AHBA gene expression dataset. Future work should move towards integrating individual-level genetic data with more comprehensive, longitudinal characterizations of lifestyle and environmental exposures.

Despite these limitations, our study provides a multi-layered perspective for understanding ethnicity/race-related influences on brain functional organization by weaving evidence from structural constraints, lifestyle factors, and transcriptomic signatures. By integrating these dimensions, we move beyond simplistic explanations of population-level brain differences and highlight the multidimensional nature of ethnicity/race. Ultimately, our findings advocate for a fundamental shift in perspective: from treating ethnicity/race as a simple causal variable to conceptualizing it as a complex construct whose effects on the brain are deeply entangled with a confluence of biological and environmental influences.

## Methods

### Datasets

This study utilized two publicly available datasets from the HCP: the HCP-YA and HCP-D datasets. The HCP-YA participants (N = 822; age: 22-37 years) were part of the HCP S1200 release[98] and matched the HCP subset used by Li et al[39]. rs-fMRI data were acquired during two sessions using a multiband sequence (voxel dimension: 2.0 mm isotropic; repetition time: 0.72 s) on a customized Siemens 3-T Skyra at Washington University in St. Louis. Each session comprised two runs with opposite phase-encoding directions, and the duration of each run was approximately 14.4 minutes. Structural images were acquired for each participant with a resolution of 0.7 mm isotropic. Family structures and self-reported ethnicity/race information were carefully considered in subsequent processing. HCP-D participants (N = 472; age: 6-21 years) were obtained from the lifespan HCP release 2.0[99]. For participants 8 years and older, rs-fMRI data were acquired during two sessions using a multiband sequence (voxel dimension: 2.0 mm isotropic) on a 3T Siemens Prisma scanner. Each session encompassed two runs with opposite phase-encoding directions, and the duration of each run was approximately 6.5 minutes. For the youngest participants (5-7 years), the total duration of rs-fMRI scanning was reduced to 21 min. Informed consent was obtained from all participants aged 18 and older; for participants younger than 18, informed permission was provided by their parents. Both the HCP-YA and HCP-D datasets were approved by the Institutional Review Board (IRB) at Washington University in St. Louis. Additional details of the data collection process and behavioural measures can be found elsewhere. All procedures in this study were conducted following the applicable guidelines and regulations, with approval from the IRB at the Beijing Institute of Technology.

### Image acquisition and processing

ICA-FIX denoised rs-fMRI data, which had undergone the minimal preprocessing steps of the HCP, were employed[100]. To improve the behavioural prediction performance, additional motion censoring and global signal regression were performed following the procedure described by Li et al[39]. Specifically, volumes with an RMS exceeding 0.2 mm were marked as censored frames. Frames immediately preceding and the two frames following these volumes, as well as any previously uncensored segments shorter than five volumes, were also marked as censored frames. Rs-fMRI runs were discarded if more than half of the frames were marked as censored. Next, the signal of the remaining uncensored frames was averaged across the cortical vertices to generate the global signal. The global signal and its first temporal derivative were regressed out from the original time courses.

### Defining individual-specific cortical parcellations and the functional connectome

The gradient MS-HBM approach was employed to define individual-specific cortical parcellations for each participant (https://github.com/ThomasYeoLab/CBIG/tree/master/stable_projects/brain_parcellation/Kong2022_ArealMSHBM)[40]. The steps of the MS-HBM approach are as follows: (i) The Connectome Workbench (https://github.com/Washington-University/workbench) is used to compute each individual’s diffusion embedding matrices for gradients; (ii) connectivity profiles for each seed region, initialization parameters and a probability spatial mask are generated; (iii) group-level priors are estimated for each ethnicity/race group on the basis of the Schaefer400 parcellations[41] and 40 independent participants; and (iv) Bayesian model estimation is performed for the generation of individual-level parcellations. The mathematical details of the MS-HBM approach are described elsewhere[40]. Both individualized and Schaefer400 parcellations were then used to compute the RSFC matrices. The vertex-wise time series, excluding the censored frames, were averaged across all the regions of interest (ROIs). For each run, the RSFC matrix was obtained across 400 regions using Pearson’s correlation analysis. The RSFC matrices were then Fisher *z*-transformed and averaged across runs, yielding two final 400 × 400 RSFC matrices for each participant.

### Predictive modelling

Predictive modelling was based on two distinct cross-validation procedures. For the ethnicity/race prediction, AA participants within each of the HCP-YA and HCP-D datasets were randomly split into 5 folds, and participants from the same family were kept together in the same fold. Hungarian matching was then applied to select demographically matched AA–WA pairs based on age, sex and RMS within each fold[12, 39]. For the lifestyle factor prediction, all HCP-YA participants were randomly split into 15 folds, with the family structure was taken care of. To ensure the generalizability of our findings, the entire data splitting and matching process for both tasks was repeated 40 times.

Following these data preparation steps, we applied 5-fold cross-validated KRR models to predict ethnicity/race and 15-fold cross-validated KRR models to predict various lifestyle factors (**Supplementary Table 1**). KRR models perform the prediction for test participants by leveraging the similarity of their RSFC patterns to those of the training participants. Specifically, the prediction for a given test participant was calculated as the weighted average of the behavioural scores from all training participants, where the weights were determined by the Pearson’s correlation between the vectorized RSFC matrices of the test participants and each training participant. For each test fold, kernel regression parameters were estimated using data from the four training folds. The optimal *l2* regularization parameter (λ) was selected through an inner 5-fold (or 15-fold) cross-validation performed exclusively on the training data. All the determined parameters were subsequently used to make predictions for participants in the test set. To mitigate the potential sensitivity to data splitting variations, the entire cross-validation process was repeated 40 times. Additionally, for each lifestyle factor, a composite score was generated before model training. This was achieved by *Z-*normalizing all related measures across participants and then averaging them for each individual. When variables other than education and income were predicted, age, sex, RMS, ICV, education, and income were included as covariates. In contrast, when predicting education and income, only age, sex, RMS, and ICV were included as covariates and regressed out from the RSFC.

For the ethnicity/race prediction models, accuracy of each test set was defined as the proportion of correctly predicted samples to the total number of samples. For the lifestyle factor prediction models, the accuracy of each test set was determined using Pearson’s correlation between the predicted and actual scores. The accuracy measures were averaged across all the folds for each data split, yielding 40 final prediction accuracy values. To evaluate the prediction significance, a corresponding set of null models was generated for each predictive model. Specifically, the output variable was permuted 1,000 times. For each permutation, a null model was trained and tested following the same procedures applied to the original model. The significance (*P* value) of each model was defined as the proportion of null models whose prediction accuracies exceed those of the true models.

### Predictive feature weights

The Haufe transformation was used to interpret the brain–behaviour relationships learned by the prediction models[46]. For each training fold, Haufe-transformed feature weights were calculated as the covariance between the input features (*RSFC*_train_) and the output predicted scores (*y’*_train_). The feature weight matrices were averaged across the 40 splits to obtain the final importance measures (400 × 400), reflecting the contribution of each edge to the prediction. We subsequently summarized the pairwise Haufe-transformed weights to the network level (17 × 17). In addition, regional importance was calculated by averaging these weights for each of the 400 regions (400 × 1). In this study, we preserved the original positive and negative signs of the feature weights rather than taking their absolute values[101].

### Morphometric similarity analysis

The Scharfer400 parcellation was transformed into the T1w surface space of each participant, and five morphometric features were extracted from the participant’s T1w images for each region, including cortical thickness, gray matter volume, surface area, Gaussian curvature and mean curvature[32]. For each participant, each morphometric feature was subjected to *Z*-normalization across regions, and a feature vector (5 × 1) was created for each region. Pearson’s correlation analysis was then employed to estimate the morphometric similarity between cortical regions, yielding a pairwise MSN (400 × 400) for each participant. The full MSN matrices were then averaged across rows to generate a regional MSN map.

To visualize the spatial pattern of the MSN across the cortex, the regional MSN maps were averaged across participants within each ethnicity/race group. To examine the ethnicity/race-related variations in the MSN, an LRM was fitted for each region. These models used the regional MSN value as the response variable and included age, sex, RMS, ICV, education and income as covariates. Weights were applied to account for differences in group sizes, with each participant assigned a weight equal to the reciprocal of the size of their corresponding group. Two-sided *t* statistics were extracted from the LRMs, and significance was determined at *P* < 0.05 with the application of FDR correction for multiple comparisons across the 400 regions to control for Type I errors.

### Structural equation modelling

The *lavaan* package (version 0.6.18) in R was used to fit a SEM for each lifestyle factor and estimate the regression coefficients for each path while controlling for age, sex, RMS and ICV[54, 55]. The complete model equations are provided in the Supplementary Materials. The latent variable for brain function was derived from the participants’ principal functional gradient. In this study, we applied principal component analysis (PCA) to the RSFC matrix of each participant for dimensionality reduction, and the resulting first principal component was identified as the principal functional gradient. The latent variable for brain function was subsequently constructed from the *Z*-normalized gradient values of the top 15 cortical regions that were most significantly correlated with each lifestyle factor[54]. Before being entered into the model, we subjected the RMS, ICV and lifestyle factors to *Z*-normalization to avoid scale effects and facilitate coefficient comparisons across models. The *P* value for each path was calculated by dividing the estimated parameter by its robust standard error (SE) and then testing the null hypothesis that the parameter equals zero. The *P* values of all paths within each SEM were then subjected to FDR correction to account for multiple comparisons. We further calculated the mediation effects (a*b) of each lifestyle factor and the proportion of ethnicity/race-related variability mediated by the lifestyle factors (a*b/(a*b+c)). FDR correction was further controlled at 0.05 across all the mediation effects, direct effects and total effects tested.

### Interaction effects modelling

To examine how the ethnicity/race-related variability in brain connectivity is moderated by the education level and substance use status of participants, we fitted two LRMs for each region. These models controlled for age, sex, RMS and ICV and included an interaction term between ethnicity/race and either education or substance use[102]. The response terms of these models were the regional principal functional gradient value, defined as the principal components derived from the RSFC matrix dimensionality reduction using PCA. Moreover, two-sided *t* statistics for the interaction terms were extracted from the LRMs, and significance was determined at *P* < 0.05, with the application of FDR correction for multiple comparisons across 400 regions to prevent Type I errors. The complete model equations are provided in the Supplementary Materials.

### Transcriptomic analysis

We utilized gene expression data from the AHBA dataset, derived from six postmortem brains (age: 24 to 57 years; 1 female) and comprising 3702 spatially distinct samples. The preprocessing steps were performed according to the approach of Markello et al.[103]. These steps are as follows: (i) probe-to-gene annotations were first updated, followed by intensity-based filtering with a threshold of 0.5 to select probes with the most consistent patterns of regional variation across donors, after which probes were aggregated to genes; (ii) gene expression samples were mapped to the Schaefer400 parcellation, in which a bidirectional mirroring approach across the left and right hemispheres was applied, and centroid-based imputation was employed to address missing data; (iii) the expression values of each sample were normalized across the genes for each donor, and subsequently, the expression values of each gene were normalized across all samples for each donor; and (iv) the samples were aggregated across donors to generate expression values for a given region. Thus, 15,632 genes remained, and a gene expression matrix (400 × 15,632) was used for subsequent analyses[104].

PLS regression was employed to relate the Haufe-transformed ethnicity/race predictive weights to the transcriptomic measurements of all 15,632 genes[32]. In the PLS model, *Z*-normalized gene expression data served as the predictor variable, and *Z*-normalized predictive weights served as the response variables. PLS1 provides the optimal low-dimensional representation of gene expression profiles that co-vary most strongly with ethnicity/race-related variability in brain functional connectivity. The statistical significance of the variance explained by PLS1 was assessed by spatially permuting the response variables 1,000 times. Finally, bootstrapping was used to quantify the estimation errors for the PLS1 weight of each gene. A *Z* score for each gene was then calculated by dividing its weight by its bootstrap SE, enabling the ranking of genes based on their PLS1 contributions.

In both the HCP-YA and HCP-D datasets, the top 2,000 positive genes and 2,000 negative genes were selected based on their *Z*-score values to form the PLS1+ and PLS1-gene sets, respectively. Each gene set was then submitted separately to the Metascape website, an automated functional pathway enrichment tool that provides biological insights for the given gene list. To validate the enrichment results, the PLS genes from the HCP-YA and HCP-D datasets were then entered into Metascape together to conduct a multi-gene list meta-analysis. This approach facilitates the understanding of pathways that are shared across gene lists or are unique to a specific gene list. All identified pathways were considered significant at *P* < 0.05, corrected for multiple comparisons by the FDR.

### Statistical analysis

The spatial association between different cortical maps was assessed across parcels using two-sided Pearson’s correlation analysis. To account for distance-dependent spatial autocorrelation, which can lead to an overestimation of statistical significance, we implemented a spin test. Specifically, a null distribution was generated for each cortical feature by performing 1,000 random rotations of the corresponding feature map. Permutation *P* values were calculated by counting the number of times that the null correlation exceeded (for positive correlations) or fell below (for negative correlations) the observed correlation, divided by the total number of spatial rotations. Spin tests were implemented using the Neuromaps toolbox (https://github.com/netneurolab/neuromaps).

## Data availability

The raw and preprocessed data are available from the HCP (https://www.humanconnectome.org/study/hcp-young-adult/document/1200-subjects-data-release; https://humanconnectome.org/study/hcp-lifespan-development). The subject lists for both the HCP-YA and HCP-D datasets are available on our GitHub page (https://github.com/TianyiYanLab/Ethnicity_Race_Diversity). Human gene expression data are available from the Allen Brain Atlas (https://human.brainmap.org). In our analyses, we also utilized publicly available cortical atlases, including the Schaefer400 atlas (https://github.com/ThomasYeoLab/CBIG/tree/master/stable_projects/brain_parcellation/Schaefer20 18_LocalGlobal/Parcellations), the S–A axis (https://github.com/PennLINC/S-A_ArchetypalAxis), and environmental effect maps (https://doi.org/10.5281/zenodo.7606653). The PLS1 gene lists and *Z* score weights for both cohorts are provided in Supplementary Data 1. All data supporting the findings of this study are included in the paper and its Supplementary Information, and all additional information is available from the authors upon reasonable request.

## Code availability

All the codes used to generate the results are available on GitHub (https://github.com/TianyiYanLab/Ethnicity_Race_Diversity).

## Acknowledgements

The authors thank the HCP team and Dr. Zaixu Cui for their support and help. This work was supported by the National Natural Science Foundation of China (grant numbers 62336002, 62406025 and 82302175); the STI 2030-Major Projects (grant number 2022ZD0208500); the Key-Area Research and Development Program of Guangdong Province (grant number 2023B0303030002); the National Science and Technology Innovation 2030 Program (grant number 2021ZD0200500); and the Beijing Nova Program (grant number 20230484465).

## Notes

https://www.humanconnectome.org/study/hcp-young-adult/document/1200-subjects-data-release

https://humanconnectome.org/study/hcp-lifespan-development

### Competing Interest Statement

The authors have declared no competing interest.

### Summary of Updates

We have revised the manuscript to enhance clarity and readability, ensuring that the presentation of results and interpretations is easier to follow. No changes were made to the scientific content or conclusions.

