## Supplementary Information for "Lifestyle and transcriptional signatures associated with ethnicity/race-related variations in the functional connectome"

**Supplementary Figure 1**

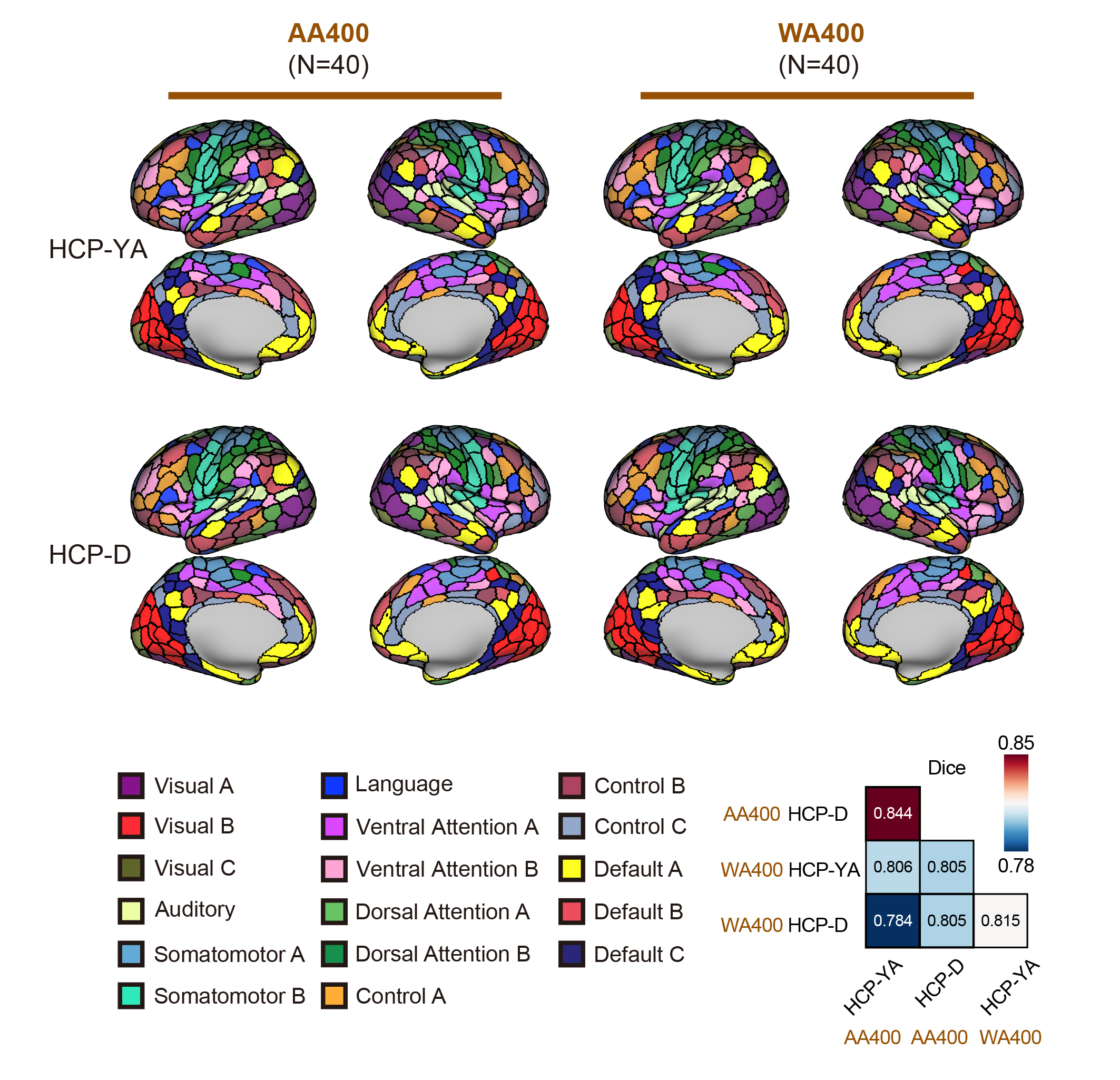

**Fig. S1 Ethnicity/race-related variability in the spatial topography of functional brain organization.** Group-level areal parcellation maps (AA400 and WA400) were generated from 40 independent AA and WA individuals’ individualized parcellations, respectively, by assigning each cortical vertex to its most likely parcel. The group-level AA400 and WA400 parcellations capture both shared and unique topographical features. Notably, parcellation maps within the same ethnic/racial group across datasets showed high similarity (Dice = 0.844 for AA; 0.815 for WA), whereas the similarity between AA and WA maps across datasets was substantially lower (Dice = 0.784).

**Supplementary Figure 2**

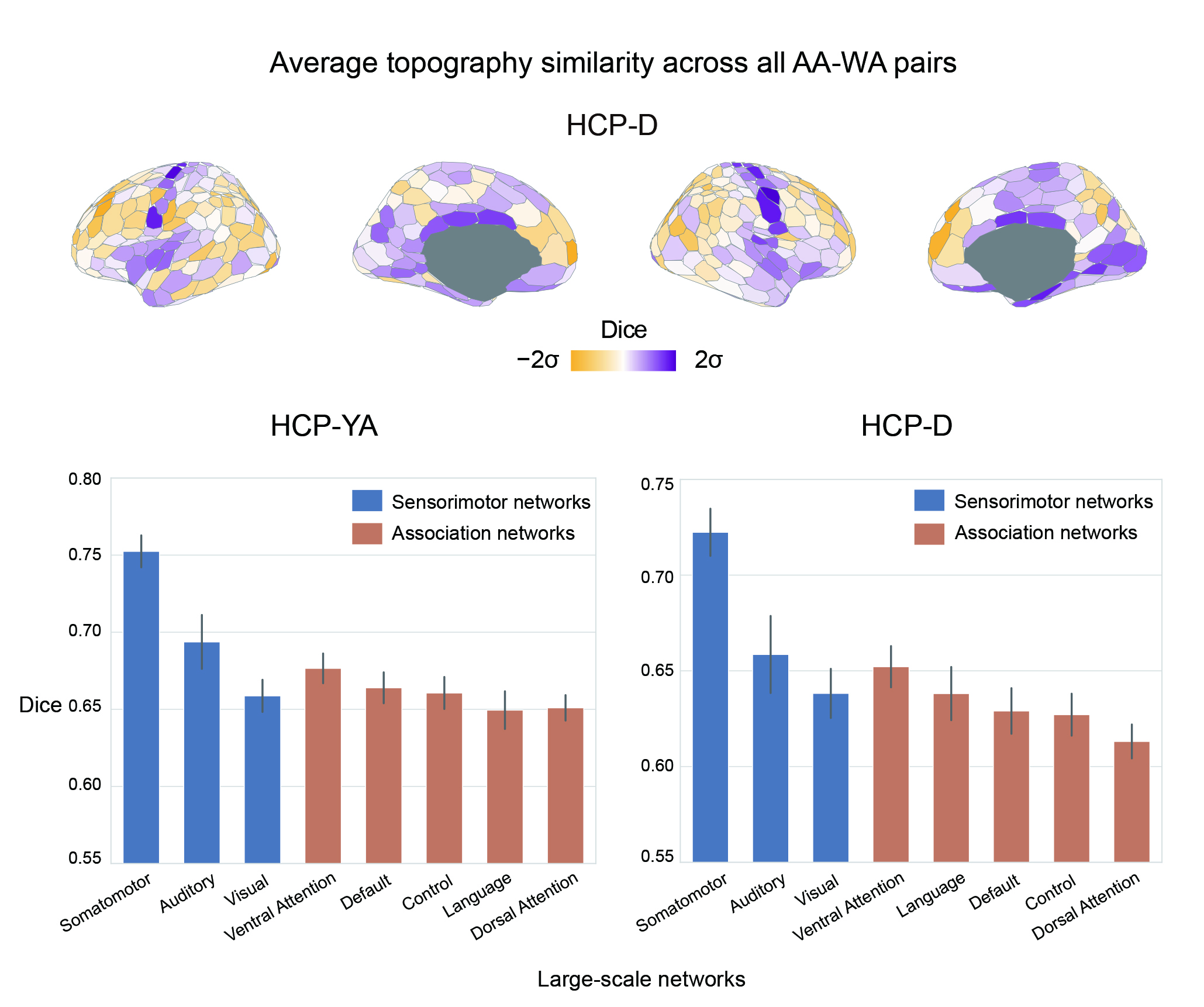

**Fig. S2** **Topographic variability of individual-specific parcellations across AA–WA pairs is higher in association cortices than in sensorimotor cortices.** Ethnicity/race-related topographic variability was more pronounced in association cortices than in unimodal sensorimotor cortices. The Yeo atlas[1] is divided into association cortical networks, including default, control, dorsal attention, ventral attention and language networks, and sensorimotor cortical networks, including auditory, somatomotor and visual networks. Each bar shows the average Dice value across regions within a network, with error bars representing the standard error of the mean.

**Supplementary Figure 3**

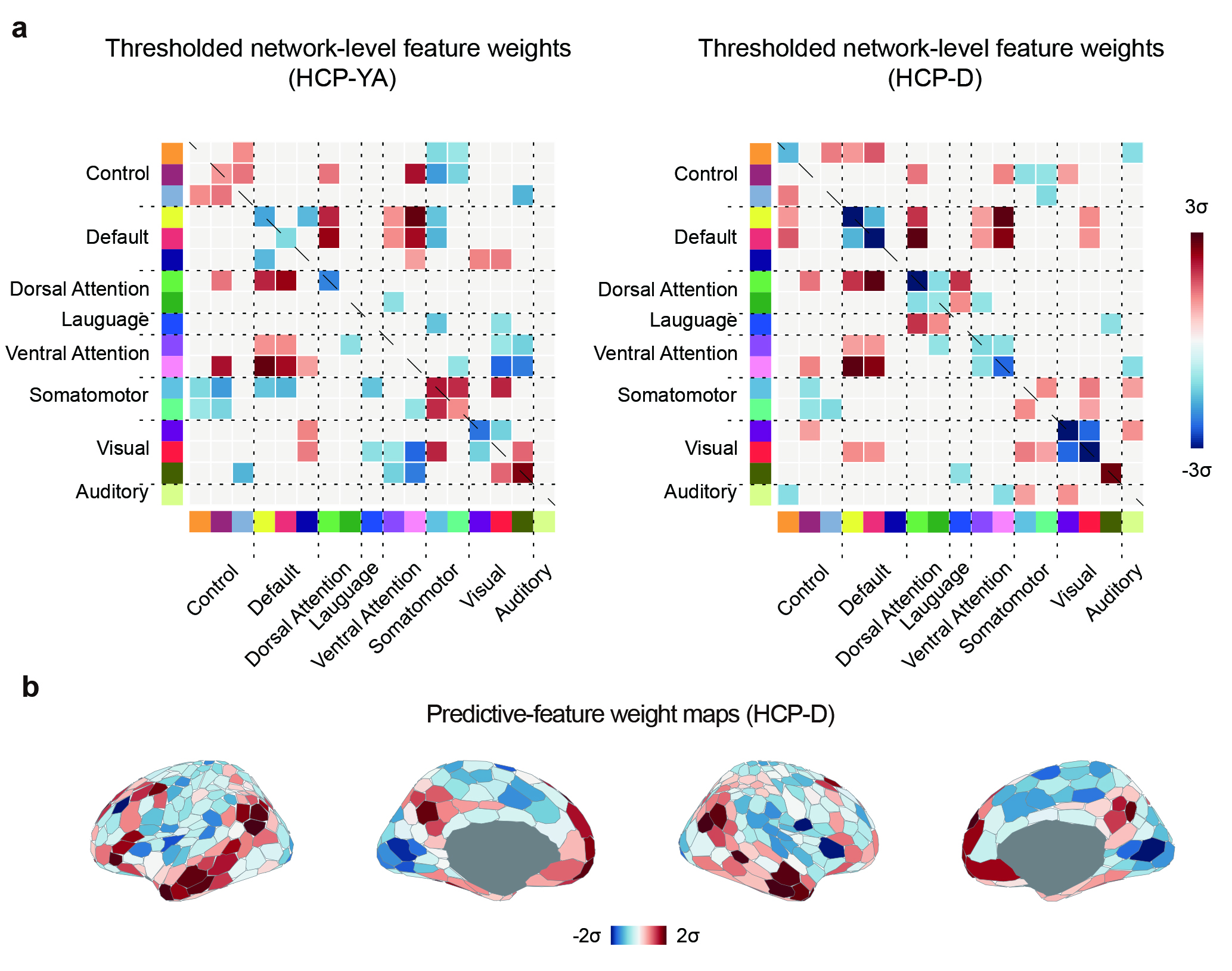

**Fig. S3 Haufe-transformed weights of the ethnicity/race prediction model.** a, Thresholded Haufe-transformed predictive feature matrices for both datasets. The top quartile of the absolute weight values from the original matrices was retained, and the corresponding original values was displayed. b, The Haufe-transformed weights of each region were averaged to generate ethnicity/race-predictive weight maps in the HCP-D dataset.

**Supplementary Figure 4**

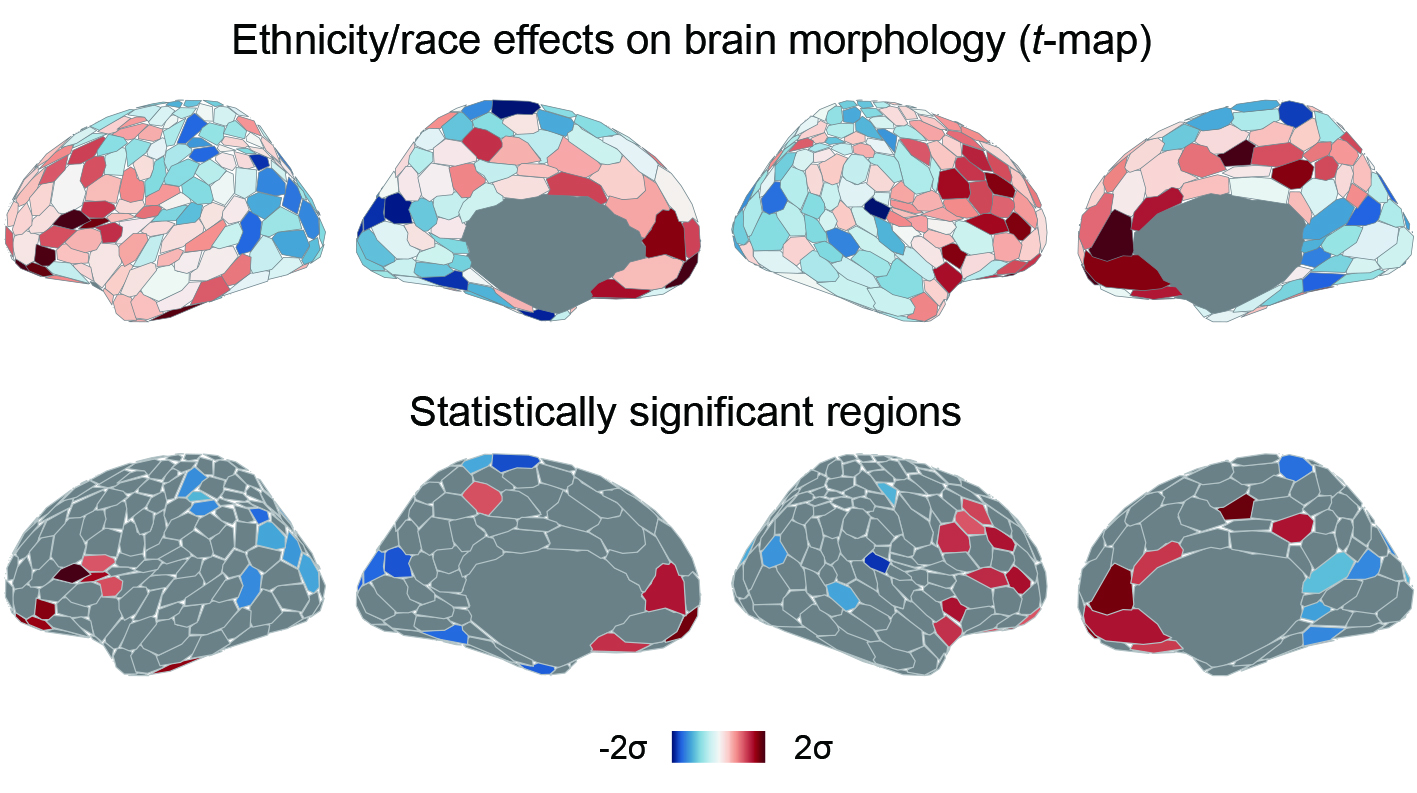

**Fig. S4** **Ethnicity/race-related variability in brain morphometric similarity patterns in the HCP-D dataset.** Top panel: Comparison (*t*-map) of regional morphometric similarity patterns across ethnic/racial groups. Bottom panel: False discovery rate (FDR) correction with *P* < 0.05 for multiple comparisons across 400 regions revealed statistically significant ethnicity/race-related effects in cortical morphometry.

**Supplementary Figure 5**

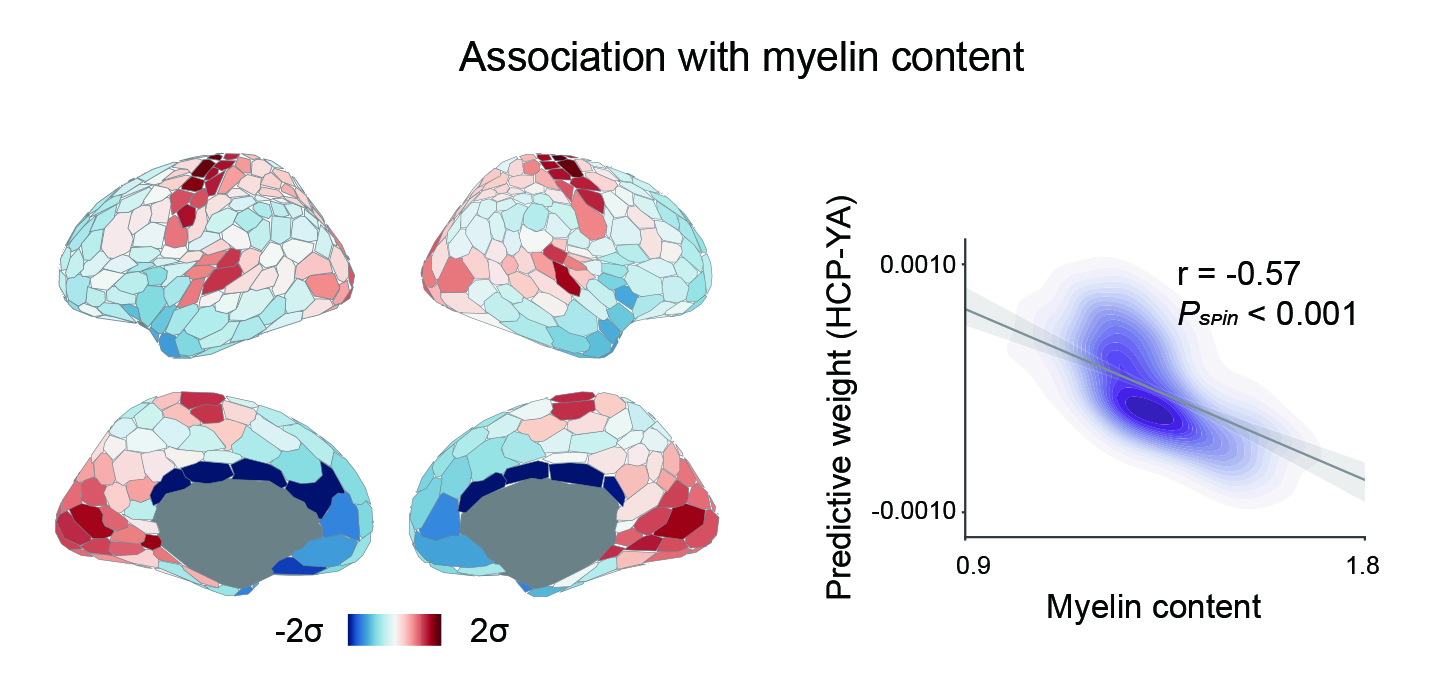

**Fig. S5** **Association with cortical myelin content.** Ethnicity/race-predictive weights were negatively correlated with the distribution of cortical myelin content (r = -0.57, *P*_spin_ < 0.001), which was estimated from the T1w/T2w ratio.

**Supplementary Figure 6**

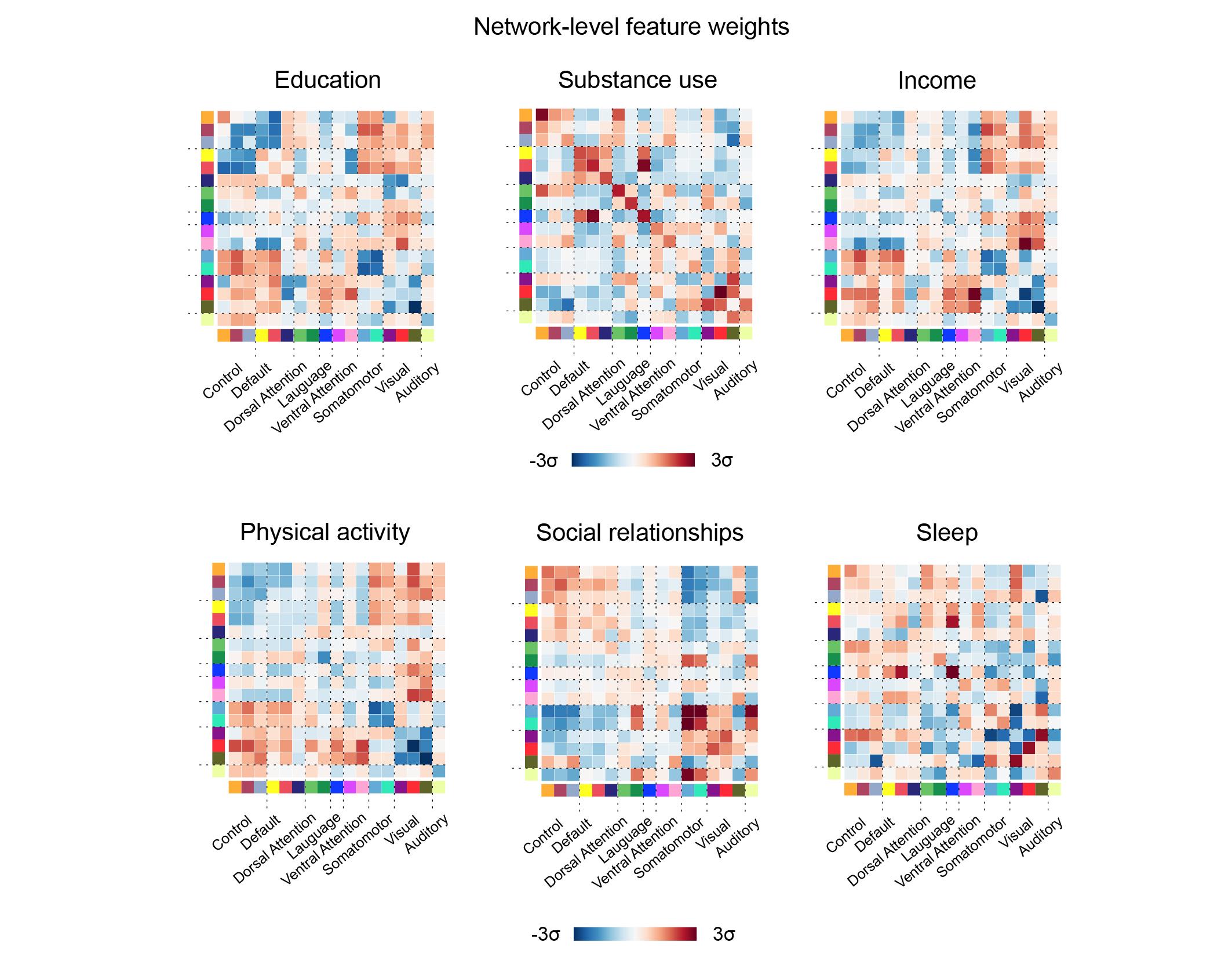

**Fig. S6 Network-level Haufe-transformed predictive feature matrices for education, substance use, income, physical activity, social relationships and sleep health measures in the HCP-YA dataset.** Regional pairwise feature weights were averaged to the network level based on the Yeo networks. For visualization, actual values were subjected to *Z* score standardization.

**Supplementary Figure 7**

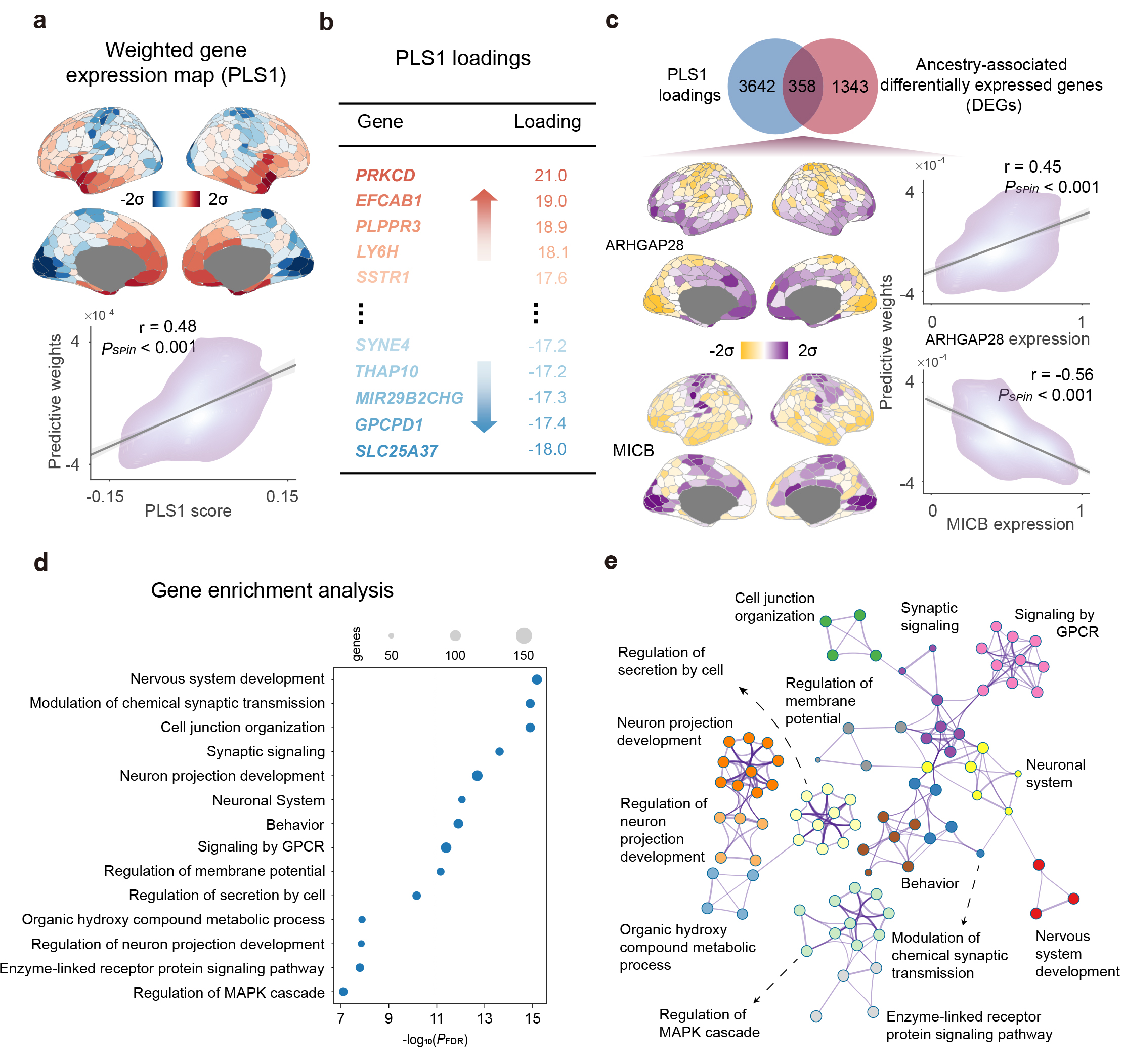

**Fig. S7** **Gene expression profiles associated with ethnicity/race-related functional connectivity variability in the HCP-D dataset**. **a,** Weighted cortical gene expression map of the regional PLS1 scores (top panel). Scatterplot showing the relationship between the PLS map and the Haufe-transformed predictive weight map (bottom panel, *r* = 0.48, *P_spin_* < 0.001). **b,** Genes were ranked according to their loadings on PLS1. **c,** The PLS+ genes overlapped with ancestry-associated differentially expressed genes (DEGs) reported by Benjamin et al.[2]. Among the 358 overlapped genes, ARHGAP28 exhibited the strongest positive association with the Haufe-transformed predictive weights (*r* = 0.45, *P_spin_* < 0.001), whereas MICB showed the strongest negative association (*r* = -0.56, *P_spin_* < 0.001). **d,** Representative enriched terms of PLS1+ genes. Circle size represents the number of genes within a given ontology term, and the multi-test FDR-adjusted *P* values are plotted as log10-transformed values. **e,** Metascape network plot of enriched terms, capturing intra-cluster and inter-cluster similarity relationships. Each term is represented by a circle, coloured by its cluster identity and scaled proportional to the number of genes involved.

**Supplementary Figure 8**

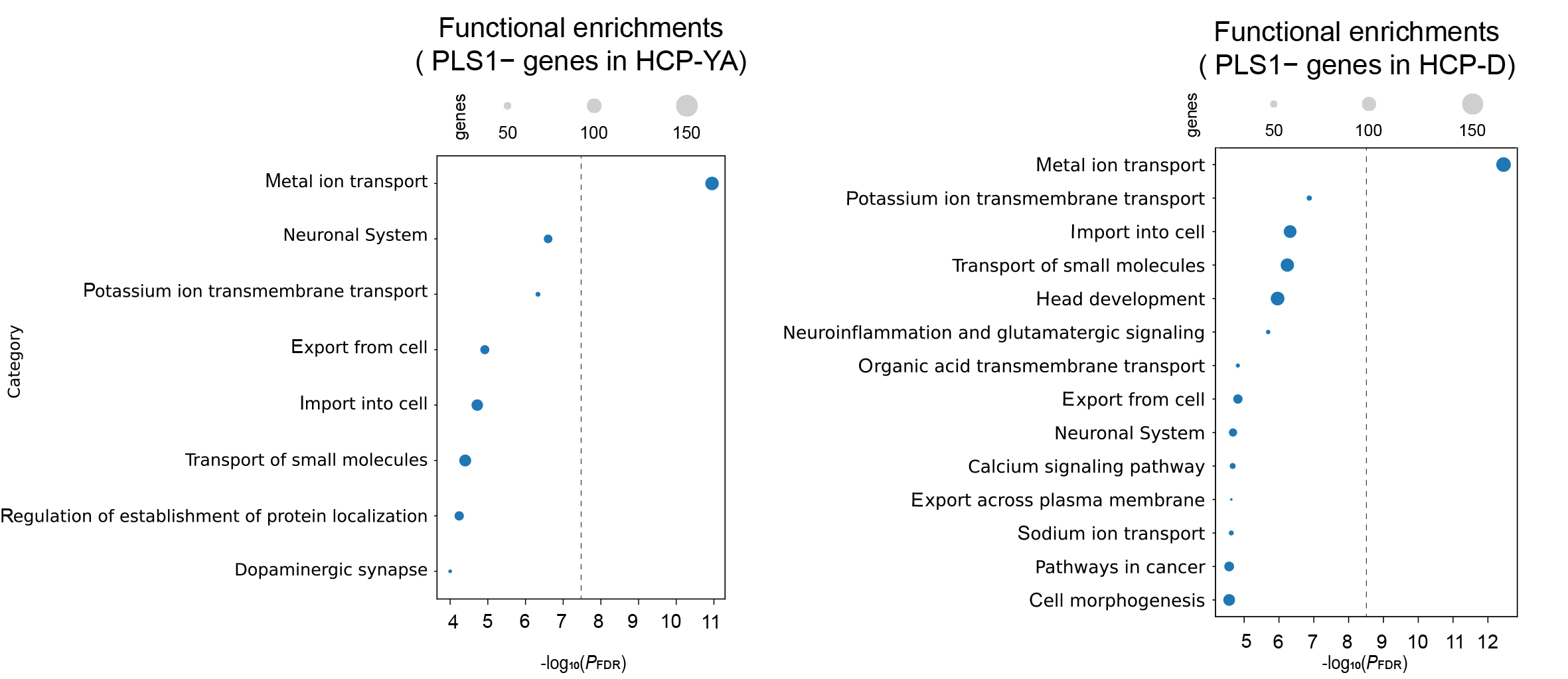

**Fig. S8 Representative enriched terms of PLS1- genes in the HCP-YA and HCP-D datasets.** Circle size indicates the number of genes within a given ontology term, and the multi-test FDR-adjusted *P* values are plotted as log10-transformed values.

**Supplementary Figure 9**

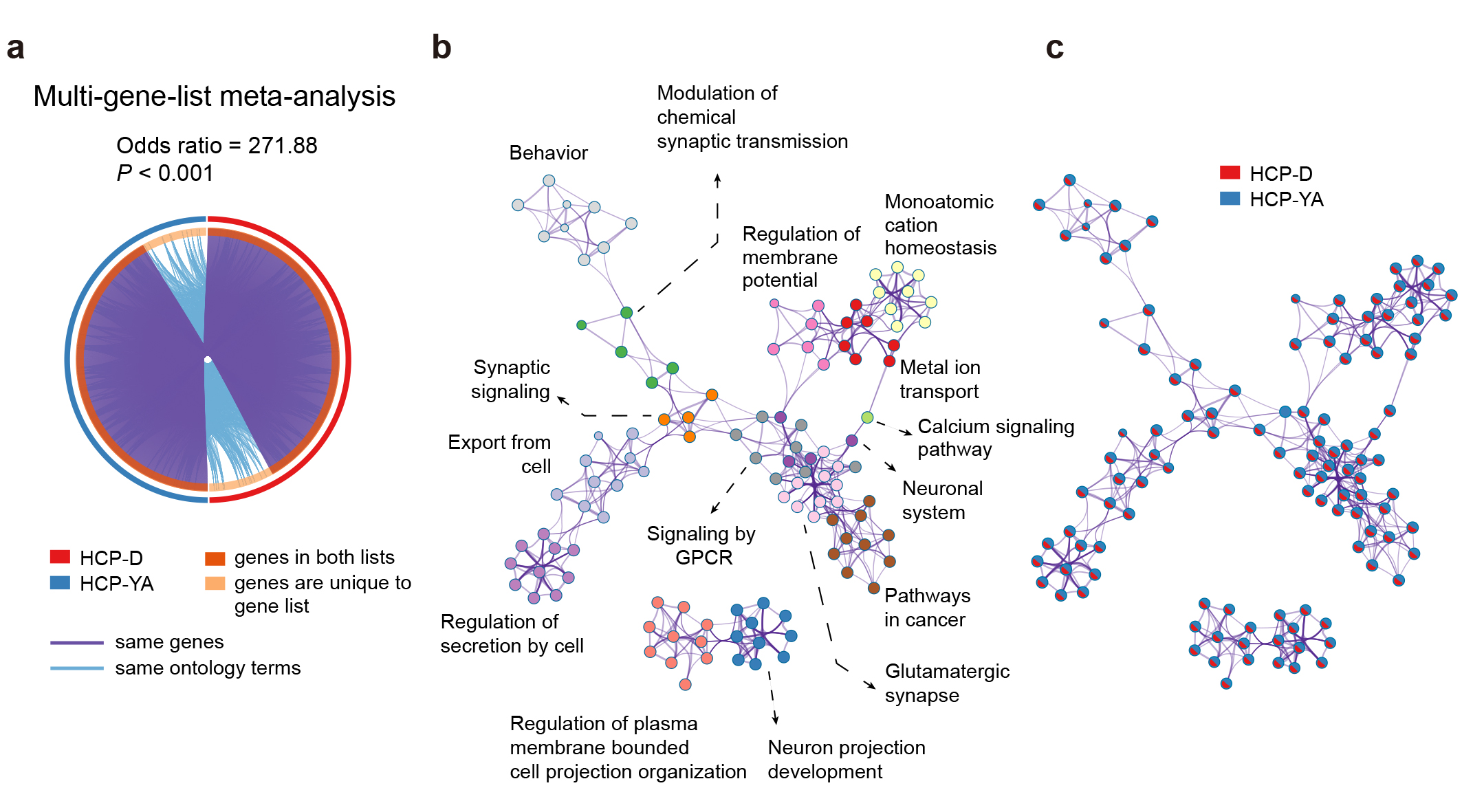

**Fig. S9 Validation of transcriptional enrichment via multi-gene list meta-analysis.** **a,** Circos plot visualizing the overlap of genes between the HCP-YA and HCP-D datasets. Purple curves connect identical genes and blue curves connect genes belonging to the same ontology term. The PLS gene lists from the two independent datasets are highly reproducible. **b,** Metascape network plot of the representative enriched terms. Each term is represented by a circle, coloured by its cluster identity and scaled proportional to the number of genes involved. **c,** The same network plot as in (b), but with the nodes represented as pie charts. The pies are colour-coded according to their corresponding gene list identities.

**Supplementary Table1 Lifestyle characteristics of participants.**

| **Behavioral domain (Dataset)** | **Formal Name** | **Intuitive Name** |
| --- | --- | --- |
| **Education** | SSAGA_Educ | Education |
| **Income** | SSAGA_Income | Income |
| **Sleep** | PSQI_Score | PSQI |
| **Substance Use** | Num_Days_Drank_7days  SSAGA_Alc_D4_Dp_Sx  SSAGA_Alc_D4_Ab_Dx  SSAGA_Alc_D4_Ab_Sx  SSAGA_Alc_D4_Ab_Sx  Num_Days_Used_Any_Tobacco_7days  SSAGA_TB_Smoking_History  SSAGA_TB_Still_Smoking  SSAGA_Times_Used_Illicits  SSAGA_Times_Used_Cocaine  SSAGA_Times_Used_Hallucinogens  SSAGA_Times_Used_Opiates  SSAGA_Times_Used_Sedatives  SSAGA_Times_Used_Stimulants  SSAGA_Mj_Use  SSAGA_Mj_Times_Used | Alcohol Use  Alcohol Dependence (Sx)  Alcohol Abuse (Dx)  Alcohol Abuse (Sx)  Alcohol Dependence (Dx)  Tobacco Use  Smoking History  Current Smoking  Illicits Use  Cocaine  Hallucinogens  Opiates  Sedatives  Stimulants  Marijuana History  Marijuana |
| **Physical Activity/Motor** | Endurance_Unadj  GaitSpeed_Comp  Dexterity_Unadj  Strength_Unadj | Endurance  GaitSpeed  Dexterity  Strength |
| **Social Relationship** | Friendship_Unadj  Loneliness_Unadj  PercHostil_Unadj  EmotSupp_Unadj  InstruSupp_Unadj | Friendship  Loneliness  PercHostil  EmotSupp  InstruSupp |

**Equation for structural equation models**

**Education:**
Y_brain function_ ~ age + sex + RMS + ICV + c*group + b*education;
education ~ a*group + age + sex
**Substance Use:**
Y_brain function_ ~ age + sex + RMS + ICV + education + income + c*group + b*substance use;
substance use ~ a*group + age + sex + education + income
**Income:**
Y_brain function_ ~ age + sex + RMS + ICV + c*group + b*income;
income ~ a*group + age + sex
**Physical Activity:**
Y_brain function_ ~ age + sex + RMS + ICV + education + income + c*group + b*physical activity;
physical activity ~ a*group + age + sex + education + income
**Social Relationships:**
Y_brain function_ ~ age + sex + RMS + ICV + education + income + c*group + b*social relationships;
social relationships ~ a*group + age + sex + education + income
**Sleep:**
Y_brain function_ ~ age + sex + RMS + ICV + education + income + c*group + b*sleep;
sleep ~ a*group + age + sex + education + income

**Equation for Interaction effects models**

Y_brain function_ ~ age + sex + RMS + ICV + group*education

Y_brain function_ ~ age + sex + RMS + ICV + group*substance use
